# Brain-muscle connectivity during gait: corticomuscular coherence as quantification of the cognitive reserve

**DOI:** 10.1101/2022.05.19.492238

**Authors:** L. Caffi, S. Boccia, V. Longatelli, E. Guanziroli, F. Molteni, A. Pedrocchi

**Affiliations:** NearLab, Department of Electronics, Information and Bioengineering, Politecnico di Milano, Milan, Italy; Villa Beretta Rehabilitation Center, Valduce Hospital, Costa Masnaga, Italy

**Keywords:** Brain-muscle connectivity, Cognitive reserve, Corticomuscular coherence, Electroencephalography, Electromyography, Gait

## Abstract

A detailed comprehension of the central and peripheral processes underlying walking is essential to develop effective therapeutic interventions to slow down gait decline with age, and rehabilitation strategies to maximize motor recovery for patients with damages at the central nervous system. The combined use of electromyography (EMG) and electroencephalography (EEG), in the framework of coherence analysis, has recently established for neuromotor integrity/impairment assessment. In this study, we propose corticomuscular (EEG-EMG) and inter/intramuscular (EMG-EMG) coherences as measures of the cognitive reserve, i.e., the process whereby a wider repertoire of cognitive strategies, as well as more flexible and efficient strategies, can moderate the manifestation of brain disease/damage. We recorded EEG signals from the main brain source locations and superficial EMG signals from the main leg muscles involved in gait in 16 healthy young adults (age ≤30 years) and 13 healthy elderly (age ≥65 years) during three different overground walking conditions (i.e., spontaneous walking, walking with cognitive dual-task, and walking with targets drawn on the floor). In all conditions, we calculated corticomuscular and inter/intramuscular coherences. We observed higher corticomuscular and inter/intramuscular coherences during targeted walking compared to spontaneous walking in both groups, even if the increase was greater in young people. Considering dual-task walking compared to spontaneous walking, only corticomuscular coherence in the elderly increased. These results suggest age-related differences in cognitive reserve that reflect different abilities to perform complex cognitive or motor tasks during gait. This study demonstrates the feasibility, repeatability, and effectiveness of the proposed method to investigate brain-to-muscle connectivity during different gait conditions, to study the related changes with age, and to quantify the cognitive reserve.

## 1. Introduction

Walking is a fundamental motor function in our lives. It is a complex task that requires the coordinated activation of several muscles [1] and precise control strategies to continuously maintain the correct balance [2] in an ever-changing environment. Recent studies have demonstrated that, in humans, the motor cortex is not only involved in the control of voluntary and precise movements and in high level motor planning (e.g., gait initiation, addressing obstacles, etc.), but it plays a significant role also in the control of distal leg muscles during stereotyped locomotion [1, 3]. A detailed comprehension of the processes underlying walking, including mechanisms involving central and peripheral nervous systems, is essential to understand possible damages at the central nervous system (CNS) [4, 5].

Probing neuromuscular control in humans is methodologically challenging, as requires non-invasive and portable tools [2]. In the last few years, the coherence analysis, a new bi-modal analysis, has spread among researchers and clinicians. It exploits the combined use of electroencephalography (EEG) and electromyography (EMG) to gain insights into the functional connectivity between the brain and the muscles, by co-registering and co-processing cortical activation and its motor correlates even during complex human locomotion tasks, such as walking [6]. Indeed, surface EMG and EEG are fully portable, non-invasive, easy to perform, and have a high temporal resolution [7]. Corticomuscular (EEG-EMG), intermuscular (EMG-EMG), and intramuscular (EMG-EMG) coherences have been claimed to be reliable physiological markers of the strength of the descending neural drive from the motor cortex during gait. Moreover, they are also able to reflect changes in gait function with aging and CNS disorders [4, 5, 7]. Many research groups found lower corticomuscular and inter/intramuscular coherence in older compared to young adults during different types of motor tasks: static contraction of ankle muscles [8], gait-like ankle movements [9], normal and visually-guided treadmill walking [5], and overground walking [7]. This may be related to morphological and/or functional changes that occur in the nervous system with aging, affecting the ability to produce oscillatory activity in the cortex, transmit the oscillations through the corticospinal tract, and finally synchronize motor unit firing [5]. In addition, reorganization of sensorimotor networks that occurs with age [10, 11] may also contribute to the observed age-related differences. For example, patterns of dedifferentiated neural activity or compensatory activation could alter network synchronization patterns and, thus, oscillatory corticospinal activity [5]. Corticomuscular and inter/intramuscular coherences have been correlated also to clinical measures of gait function in patient after stroke. Kitatani et al. [4] found reduced paretic side inter/intramuscular coherence compared to the non-paretic side and healthy controls that was positively correlated to gait speed. Moreover, they found significant antagonist ankle muscle coherence in patients after stroke during gait, whereas this is generally not observed in healthy subjects. This probably reflects the reorganization of the sensorimotor network aimed at compensating the reduced descending neural drive on the paretic side and the paretic plantar flexor muscle weakness to enhance stability and restore functional walking [4]. Therefore, EMG-EEG and EMG-EMG coherences have recently established in the clinical assessment of neuromotor integrity/impairment. In particular, they have been widely used to monitor the effect of healthy aging on the neuromusclular system and the recovery of motor function after stroke [12].

However, most of the works available in literature using EEG and EMG to assess brain-muscle connectivity explored either two different populations, in terms of age and/or state of health, in the same experimental condition [4], or the same population under different experimental conditions [2, 13]. Moreover, most of them recorded EEG and EMG during quasi-static tasks resembling real-world walking (e.g., cyclic ankle movements [13]), or during treadmill walking [5]. However, Roeder et al. [7] demonstrated that treadmill walking alters gait kinematics, gait performance, leg muscle activation as well as a number of spectral measures compared to overground walking. Thus, a deep understanding of corticomuscular communication during overground gait is needed.

At the same time, the coherence analysis could be exploited to quantify the “cognitive reserve” (CR). A recent study have exploited coherence analysis to inspect the relationship between age, cognitive reserve and brain connectivity (i.e., EEG-EEG coherence) [14]. The CR is referred as the process whereby a wider repertoire of cognitive strategies, as well as more flexible and efficient strategies, can moderate the manifestation of brain disease/damage [15]. High intelligence quotient, participation in leisure activities, education, and occupational level are the major components of CR [16, 17]. Early childhood factors, as childhood intelligence quotient, are highly important for the development of CR. Yet, CR is not fixed in childhood, but continues to be affected by events and circumstances as they unfold across the lifespan, such as educational attainment and occupation, that can limit the progressive reduction of CR during adult life [16, 18]. Only in the last year, the CR has been correlated with gait function, demonstrating that higher CR was associated with better neural efficiency of walking in the elderly [19]. However, the CR is currently assessed using a mixture of qualitative and quantitative measures [20]. Therefore, a reliable and standardized method for CR quantification is needed.

Within this framework, this work aims to: i) combine and further deepen the knowledge about brain-muscle connectivity during gait, ii) study the related changes with age and iii) quantify the cognitive reserve. To this aim, we performed a multi-factorial protocol that involved two different populations in term of age: i) healthy subjects with age *≤*30 years and ii) healthy subjects with age *≥*65 years, in three different overground walking conditions: i) spontaneous walking, ii) walking with cognitive dual-task and iii) walking with targets drawn on the floor. The protocol aimed at progressively shifting the subject’s cognitive functions (i.e., the CR) from the domain of cognitive activities (e.g., calculations), to the domain of motor activities (e.g., walking with precise targets). The hypothesis was that the CR should reflect the capability of the subject to translate the additional cognitive load required by the two types of tasks proposed (i.e., cognitive and motor), as captured by corticomuscular coherence, into effective recruitment, control, and coordination of leg muscles to obtain physiological gait, as captured by inter/intramuscular coherence.

## 2. Materials and methods

### 2.1. Participants

We conducted a pilot study at the Politecnico di Milano. The study was approved by its ethical committee (Ref. No. 12/2021). All subjects gave informed written consent in accordance with the Declaration of Helsinki. Participants were divided into two groups according to the age: i) healthy subjects with age ≤30 years and ii) healthy subjects with age *≥*65 years. All healthy participants were able to walk unassisted and reported no history of neurological or orthopaedic disorders. The exclusion criteria were as follows: i) body mass index <19 or >32; ii) Berg Balance Scale score <50 (out of 56); iii) Mini-Mental State Examination score <25 (out of 30); iv) use of assistive device for walking; v) self-reported difficulty in performing mobility tasks; vi) experienced a fall within the previous year; vii) involuntary weight gain or loss exceeding 5 kg within the past 6 months; viii) resting blood pressure exceeding 160/95 mmHg; ix) heart attack or symptomatic cardiovascular disease in the past year; x) orthopaedic problems affecting movement in the past year; xi) medical condition affecting movement; xii) terminal illness; xiii) cognitive disorders; xiv) history of neuromuscular disorder.

### 2.2. Experimental protocol

The designed protocol consisted of three tasks: i) spontaneous walking: the participant walked along a straight path with no obstacles; ii) dual-task walking: the participant walked along a straight path with no obstacles, and, at the same time, performed mathematical calculations loud speaking. The calculations consisted in counting backward of 7 starting from 100 [21–24]; iii) targeted walking: the participant walked along a straight path with target points to be stepped on and areas of discontinuity to avoid being drawn on the floor. Each task lasted two minutes with a resting period in between. The task execution order was randomized to remove possible undesired biasing effects on the results. Participants walked at a self-selected preferred slow walking speed.

### 2.3. Experimental setup

The muscle activity was recorded bilaterally with the BTS FREEEMG 1000 wireless electromyograph (BTS Bioengineering, Italy). Six EMG electrodes were placed over the bellies of the soleus muscle (SOL), rectus femoris muscle (RF), and semimembranosus muscle (SM) of both legs. Two electrodes were placed bilaterally over the proximal portion of the tibialis anterior muscle (TAp). The ninth electrode was placed over the distal portion of the tibialis anterior muscle (TAd) of the dominant leg. Placement of the electrodes were conducted accordingly to SENIAM (Surface Electromyography for the Non-Invasive Assessment of Muscles) guidelines [25]. Brain activity was recorded using HELMATE wireless and dry electroencephalograph (Ab Medica, Italy) with eight dry electrodes placed according to the international 10–20 system (Fp1, Fp2, Fz, C3, C4, Cz, O1, O2). Finally, to detect gait phases, we used footswitch probe (BTS Bioengineering, Italy), which allowed the definition of the foot-floor contact phases during motion, thanks to three footswitch sensors attached respectively under the tip, the metatarsal and the hell of the right foot.

### 2.4. Data analysis

#### 2.4.1 Preprocessing

EMG signal was acquired at 1000 Hz, while the EEG signal was acquired at 512 Hz. Considering EEG signal, recordings that presented excessive movement artifacts (i.e., amplitude > 400 *µ*V) occurring with regular rhythmicity in each gait cycle were excluded from subsequent analysis [7]. Then, the signals were band-pass filtered in 1-120 Hz band. EMG signals, instead, were rectified and low-pass filtered at 120 Hz. EMG and EEG signals were then time normalized.

The second part of the preprocessing was focused on the removal of motion artifacts from the EEG signals and consisted in two operations. Firstly, we applied a procedure proposed by Gwin et al. [26]. For each channel, a motion artifact template was created by averaging each step of the EEG signal. Then, the motion artifact template was fitted on each step, performing a linear operation to minimize the difference between the template and the original step. Finally, the obtained fitted artifact was subtracted from the original step signal. The operation was performed for all the steps identified in the EEG trace. Secondly, we applied Artifact Subspace Reconstruction algorithm to the data cleaned with the previous method. The cutoff parameter for determining the rejection threshold of the method was set to 10, as suggested by Chang et al. [27]. Signal processing was performed on MATLAB R2021a software (The MathWorks, Inc., USA) and with the EEGLAB Toolbox [28].

#### 2.4.2 Coherence analysis

To study brain-muscles connectivity, we calculated corticomuscular, intermuscular, and intramuscular coherences using wavelet analysis. Given a time series *x*(*t*), its continuous wavelet transform (CWT) *W*_*x*;*ψ*_, with respect to the wavelet *ψ*(*t*), is a function of a translation parameter that control the location of the wavelet in the time domain (*τ*), and a scaling factor that controls the width of the wavelet (*s*). It is computed as follows:

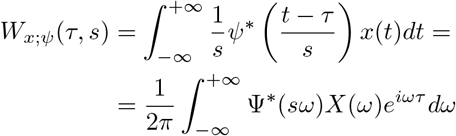

The generalized Morse wavelet was chosen, as suggested by Lilli et al. [29], and computed as:

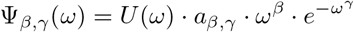

where *U* (*ω*) is the Heaviside step function, *a*_*β,γ*_ is a normalizing constant, *γ* characterizes the symmetry of the Morse wavelet and *β* · *γ* is the time-bandwidth product. The properties of Ψ_*β,γ*_(*ω*) change with *β* and *γ*. We selected *β* = 40 and *γ* = 5 following a trial and error method in order to have a balanced time and frequency resolution. Then the wavelet coherence was calculated as:

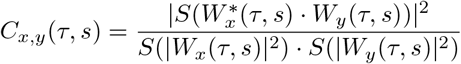

where *x* and *y* are two one-dimensional time series, *S* is the smoothing operator in time and frequency, and *W*_*x*_(*τ, s*) and *W*_*y*_(*τ, s*) are respectively the CWTs of *x* and *y* [13]. Smoothing was applied using a moving average filter with a window of 100 and 5 data points, respectively in time and frequency. Wavelet analysis was used to calculate corticomuscular coherence for each possible brain location-muscle pair, intermuscular coherence between each couple of muscles, and intramuscular coherence between right leg TAp and TAd. Afterward, each coherence was averaged on the gait cycle.

One of the most critical issue of coherence analysis is the computation of the significance threshold for the coherence itself. To this aim, for each participant, each task, and each couple of channels, we calculated the wavelet coherence also between two surrogate signals. They were obtained from the original ones with the iterative amplitude adjusted Fourier transform algorithm [30]. From the surrogate coherences, averaged on the gait cycle, we retrieved channel-specific thresholds that were used to select only the significant portions in the time-frequency domain of the coherences calculated on real data. The threshold was obtained as the median value computed pixel by pixel of all the surrogate coherences obtained for each subject and each task for the specific channel (fig. 1). To validate the generalizability of the threshold, we calculated the coefficient of variation (CV), i.e., mean/standard deviation, pixel by pixel of all the surrogate coherences calculated for that couple, considering all the subjects during the three experimental sessions. The threshold was considered reliable if the CV was lower than 20% [31]. Finally, each gait-cycle-averaged coherence and each threshold were binned across frequency and time resulting in one pixel per Hz and 1 pixel per 1% of the gait cycle.

**Figure 1.**
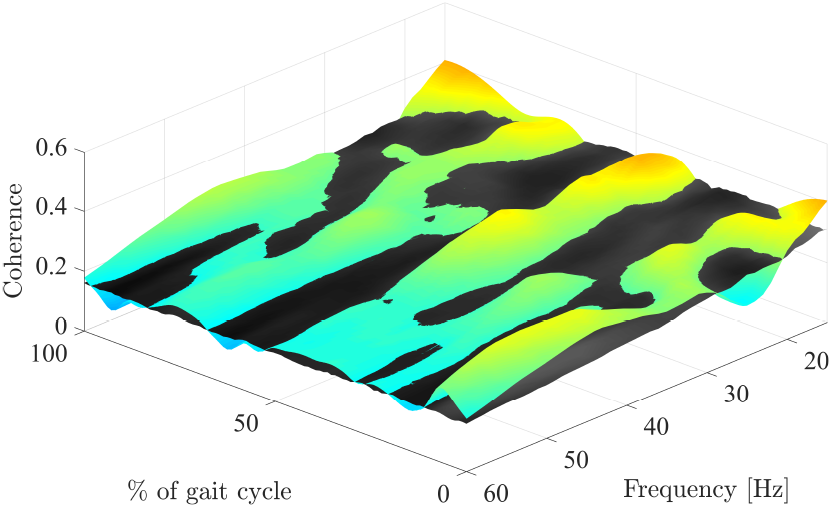
One explanatory wavelet coherence for right TAp - C3 couple (colored) and the corresponding threshold (gray).

#### 2.4.3 Outcome measures

The primary outcome measure of this work was the volume of significant coherence (i.e., volume above threshold) for each gait-cycle-averaged coherence. The volume was measured in Hz multiplied by the percentage of the gait cycle (*Hz* %*gaitcycle*). Volume above threshold analysis was conducted according to the following hypothesis:

- We considered the beta band (15-30 Hz) and the gamma band (30-60 Hz) contributions. The alpha band was not analyzed since patients after complete spinal cord injury show coherence only in this frequency band, suggesting that the origin of alpha band coherence is likely to be only spinal, thus not including the cortical contribution detected by the EEG [4].
- The phases of the gait cycle containing heel strikes were excluded from further analysis due to the presence of residual movement artifacts on EEG signals that could have altered the results, as performed by Spedden et al. [5]. Therefore, only single support phases of the gait cycle (i.e., 20%-40% and 70%-90%, as single support of the right and left leg, respectively) were retained;
- Proximal leg muscles (i.e., RF and SM) were excluded from the analysis due to possible influences of spinal column, hip, and pelvis factors, which could have hidden the task-related changes in coherence that we were interested in exploring. Therefore, only distal leg muscles (i.e., TAp, TAd, and SOL) were retained;
- Only corticomuscular coherences involving C3 and C4 electrodes were analysed due to their physiological meaning. Indeed, they are the locations most sensitive to the motor cortex activity involved in the control of the controlateral side of the body. As a consequence, C3 electrode was coupled with right distal leg muscles, while C4 electrodes with distal muscles of the left leg.

Therefore, we identified four possible time-frequency portions for coherence analysis (i.e., 20%-40% beta, 70%-90% beta, 20%-40% gamma and 70%-90% gamma) obtained by combining one phase of the gait cycle between 20%-40% and 70%-90% and one frequency band between beta and gamma. Among all the possible couples of channels, we focused on the analysis of coherence between: i) agonist and antagonist muscles of the distal legs separately (i.e., right TAp - right SOL, left TAp - left SOL), ii) two ends of the same distal leg muscle (i.e., right TAp - right TAd), iii) each distal leg muscle and the controlateral central electrode between C3 and C4 (e.g., for the right soleus muscle we considered the couple right SOL - C3).

A summary of the overall processing pipeline is represented in fig. 2. In addition, for each participant and task we calculated the median stride duration (i.e., the time interval between two right heel strikes) over all the strides detected in the recording.

**Figure 2.**
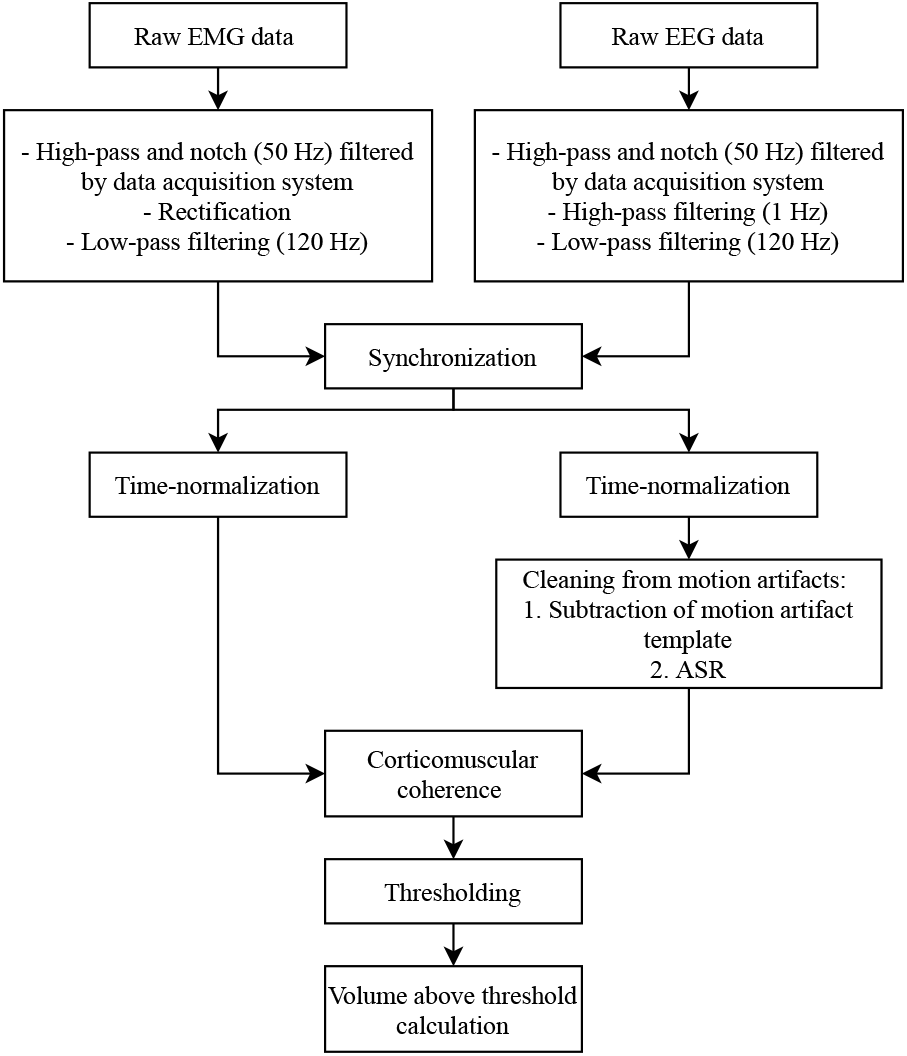
Block diagram of the processing pipeline for corticomuscular coherence.

#### 2.5. Statistical analysis

To compare the volume above threshold between tasks (i.e., spontaneous walking, dual-task walking and targeted walking) and groups (i.e., healthy young adults and healthy elderly), we implemented a generalized linear mixed model. Group and task were entered as fixed effects, task by group as interaction term, patients as random factor, and volume above threshold as dependent variable. In case of significant group or interaction effect, we performed a post-hoc analysis with the contrast on the group term to detect significant changes between groups in the three tasks separately. In case of significant task or interaction effect, instead, the post-hoc analysis was conducted with a contrast on the task term in order to inspect the differences between tasks for the two groups separately.

Moreover, median stride duration was selected as feature to compare the gait execution in the two groups during the three walking conditions. The median stride duration was compared between tasks in the two groups separately using Friedman test with post-hoc analysis conducted by means of Wilcoxon matched-pairs signed rank test. To compare the median stride duration between groups for the three tasks separately, Mann–Whitney U test was used. Analyses were performed using SPSS Statistics (Version 26) and R.

## 3. Results

### 3.1. Participants

We recruited 16 healthy young and 13 healthy elderly volunteers. The young group was composed of 9 females, 7 males, with a median (first quartile - third quartile) age of 24 (24 - 25.25) years. The elderly group, instead, was constituted by 4 females, 9 males, with median age of 69 (68 - 71) years. All participants were able to successfully complete the experimental protocol.

### 3.2. Data analysis

. We removed from the analysis a median (first quartile - third quartile) of 0 (0 - 2) channels for the young group and of 0 (0 - 1.5) for the elderly group due to excessive movement artifacts.

Concerning the validation of the thresholding method, we were able to compute 87 surrogate coherences for each couple of channels, derived from a total number of 29 participants that underwent three experimental conditions (i.e., spontaneous walking, dual-task walking, and targeted walking). For all couples, the CV was lower than 20% (the maximum median value obtained was 16.9%, see Supplementary Materials - fig. 1). Therefore, the thresholding method was considered reliable and was applied in the analysis.

### 3.3. Outcome measures

In table 1 and table 2, the results of the statistical analysis on volumes above threshold are shown (see also Supplementary Materials - table 1, fig. 2 and fig. 3). In particular, table 1 shows the main analysis together with the post-hoc performed if a significant task or interaction terms was detected. Whilst, table 2 represents the main analysis and the post-hoc conducted in case of significant group or interaction term. For each one of the considered couples, linear generalized mixed model detected a significant task or interaction effect at least in one time-frequency portion. Statistically significant difference between groups or interaction effect were detected in all muscle couples at least in three time-frequency portion, but not in all muscle-brain source location couples. Considering the first post-hoc analysis (i.e., group contrasted), focusing on the TA muscle, we found a significant difference in the corticomuscular coherence for the left leg (left TAp-C4) in the beta band for the dual-task walking (DT) and the targeted walking (TW) conditions between the two groups in the 70%-90% of the gait cycle (GC, p-value = 0.001 for DT, and p-value = 0.004 for TW). The coherence was higher in the elderly group (EG). This difference was, instead, not detected on the same analysis on the specular couple (right TAp-C3). The post-hoc analysis for corticomuscular coherence with the SOL muscle, instead, was not conducted because we did not find any difference for both legs. Considering the intermuscular coherence for the left leg (left SOL - left TAp), we found significant differences between groups in the DT for the gamma band in the 70%-90% of the GC (p-value = 0.020). The young group (YG) showed an higher coherence. Moreover, we detected significant differences also for the TW in the beta band during the 20%-40% of the GC (p-value = 0.003), and for the gamma band in both phases of the GC (p-value = 0.010 for 20%-40% of the GC, and p-value = 0.036 for the 70%-90% of the GC). In all three cases, the coherence of the YG was higher. Considering the intermuscular coherence of the right leg (right SOL - right TAp), we found significantly higher coherence in YG compared to EG in all the tasks In particular, the significant interaction was found for spontaneous walking (SW) in gamma band for 20%-40% (p-value = 0.02) and 70%-90% of the GC (p-value = 0.016); for DT in beta band for 70%-90% of the GC (p-value = 0.042) and in gamma band for both 20%-40% (p-value = 0.03) and 70%-90% (p-value = 0.012) of the GC; for TW in beta band for both 20%-40% (p-value = 0.024) and 70%-90% (p-value < 0.001) of the GC, and in gamma band for 70%-90% of the GC (p-value = 0.002). The intramuscular coherence (right TAp - right TAd) showed significant differences between groups for all the experimental conditions for the beta band during the 70%-90% of the GC (SW: p-value = 0.044, DT: p-value = 0.030; TW: p-value = 0.012). In the gamma band, we observed a significant change only in the TW for the 20%-40% of the GC (p-value = 0.047). Also in this case, the YG was characterized by a greater value of coherence

**Table 1.**
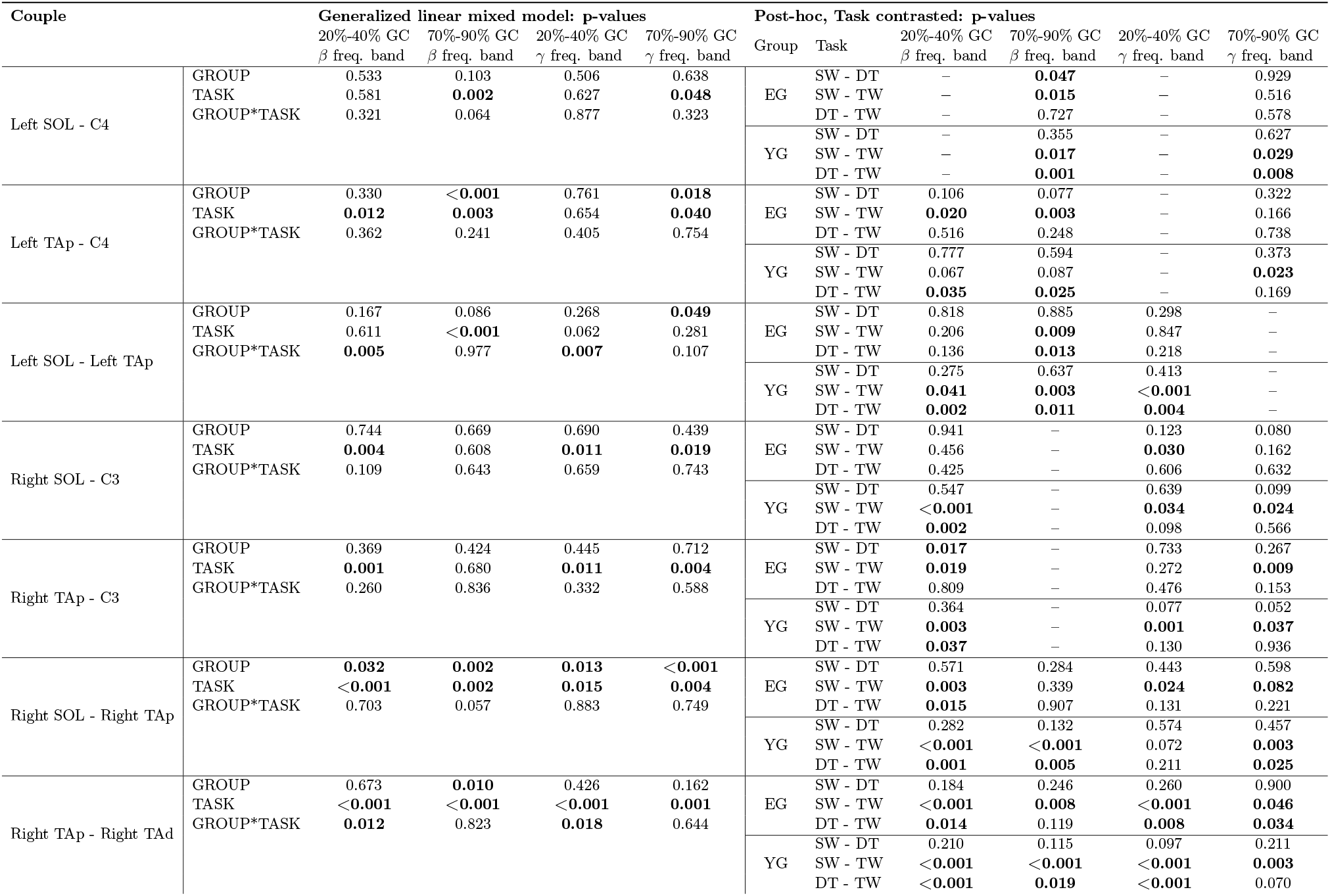
Statistical analysis to compare volume above threshold across groups (EG = elderly group and YG = young group) and tasks (SW = spontaneous walking, DT = dual-task walking and TW = targeted walking) for each couple of channels. Post-hoc analysis performed with contrast on task. The first column displays the couples considered by the statistical analysis. The central column presents the p-values of the generalized linear mixed models for group, task, and interaction term. The right column displays the post-hoc analysis with contrast on task. Results are divided for frequency bands (beta and gamma), and gait cycle (GC) phases (20%-40% and 70%-90%). Significant p-values are displayed in bold.

**Table 2.**
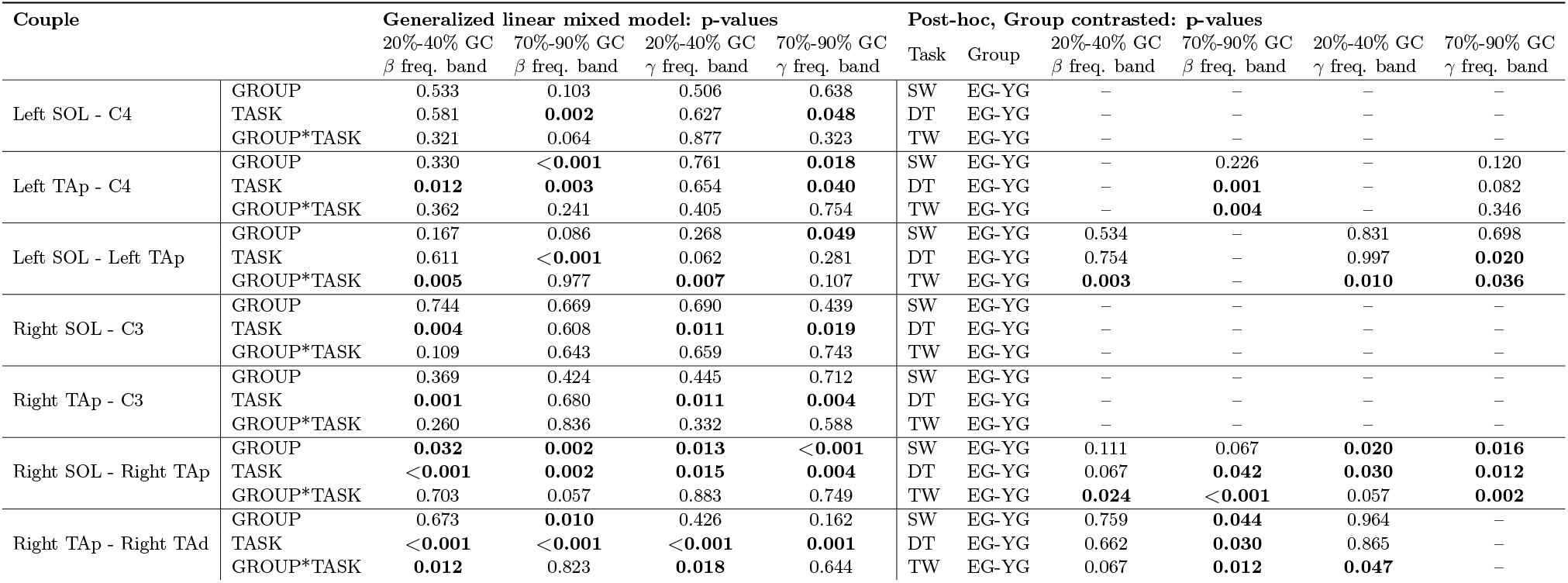
Statistical analysis to compare volume above threshold across groups (EG = elderly group and YG = young group) and tasks (SW = spontaneous walking, DT = dual-task walking and TW = targeted walking) for each couple of channels. Post-hoc analysis performed with contrast on groups. The first column displays the couples considered by the statistical analysis. The central column presents the p-values of the generalized linear mixed models for group, task, and interaction term. The right column displays the post-hoc analysis with contrast on groups. Results are divided for frequency bands (beta and gamma), and gait cycle (GC) phases (20%-40% and 70%-90%). Significant p-values are displayed in bold.

**Figure 3.**
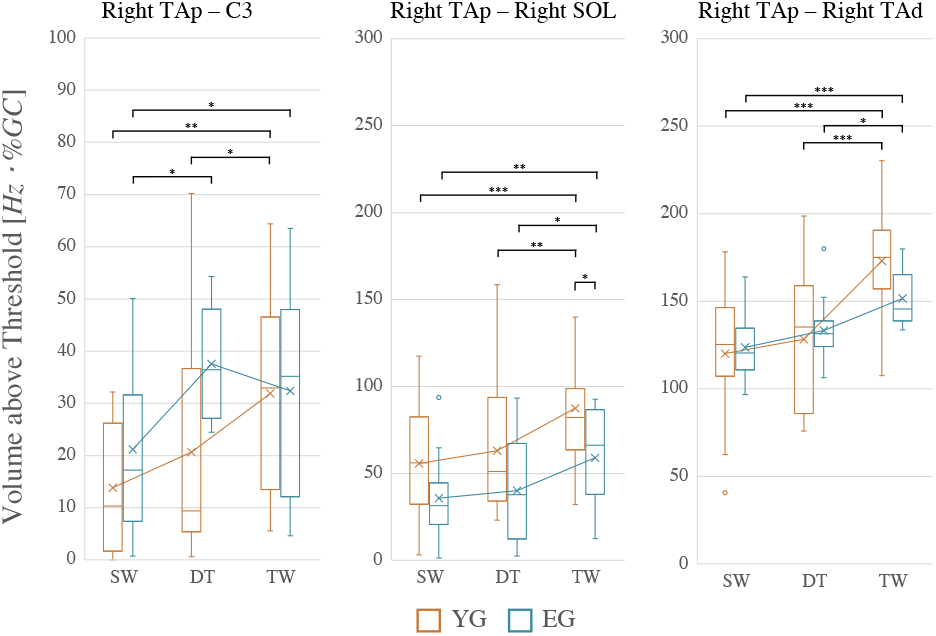
Volume above threshold in beta band and in 20%-40% of the gait cycle for one corticomuscolar, one intermus-cular and one intramuscular couple for the two groups of subjects (YG = young group and EG = elderly group) during the different walking conditions (SW = spontaneous walking, DT = dual-task walking and TW = targeted walking).

Considering the second post-hoc analysis (i.e., task contrasted), focusing on the TA muscle, we found a significant difference in the corticomuscular coherence for the left leg (left TAp - C4) for EG between the SW and TW in beta band both in the 20%-40% (p-value = 0.020) and 70%-90% of the GC (p-value = 0.003). The coherence was higher in the TW. In gamma band was detected a significant difference for YG between SW and TW (p-value = 0.023). The coherence was higher in TW.

The specular couple (right TAp - C3) presented significant differences between SW and TW for EG in beta band for 20%-40% of the GC (p-value = 0.019) and gamma band for 70%-90% of the GC (p-value = 0.009), while for YG in beta and gamma band for 20%-40% of the GC (respectively p-value = 0.003 and p-value = 0.001) and in gamma band for 70%-90% of the GC (p-value = 0.037). Coherence in TW was always higher compared to SW. EG showed also higher coherence in DT compared to SW in beta band 20%-40% of the GC (p-value = 0.017). Focusing on SOL muscle, we found significant difference in the corticomuscular coherence for the left leg (left SOL - C4) between SW and TW both in EG (p-value = 0.015) and in YG (p-value = 0.017) in beta band for 70%-90% of the GC. The same interaction was found for YG in gamma band for 70%-90% of the GC (p-value = 0.029), but not in EG. In all cases, coherence during TW was higher compared to SW. In addition, EG showed higher coherence in DT compared to SW in beta band for 70%-90% of the GC (p-value = 0.047). The specular couple (right SOL - C3) displayed significant differences between SW and TW for EG in gamma band for 20%-40% of the GC (p-value = 0.03), while for YG in beta band for 20%-40% of the GC (p-value < 0.001) and in gamma band both for 20%-40% of the GC (p-value = 0.034) and for 70%-90% of the GC (p-value = 0.024). In all cases, coherence during TW was higher compared to SW. Considering the intermuscular coherence for the left leg (left SOL - left TAp) EG presented changes in coherence between SW and TW in beta band for 70%-90% of the GC (p-value = 0.009). The same interaction was found in YG but in different time-frequency portions: in beta band for 20%-40% of the GC (p-value = 0.041), in beta band for 70%-90% of the GC (p-value = 0.003) and in gamma band for 20%-40% of the GC (p-value < 0.001). Both in EG and YG coherence was higher in TW compared to SW. Considering the intermuscular coherence of the right leg (right SOL - right TAp), we found significant changes between SW and TW in beta band for 20%-40% of the GC both in EG (p-value = 0.003) and YG (p-value < 0.001), while only in YG for 70%-90% of the GC (p-value < 0.001). For the gamma band instead the interaction was significant in 20%-40% of the GC only for EG (p-value = 0.024) and both for EG and YG in 70%-90% of the GC (respectively p-value = 0.082 and p-value = 0.003). In all cases coherence during TW was higher compared to SW. Finally, the intramuscular coherence (right TAp - right TAd) showed significant differences between SW and TW for all time-frequency locations both in EG and YG (in beta band for 20%-40% of the GC: p-value < 0.001 both for EG and YG; in beta band for 70%-90% of the GC: p-value = 0.008 for EG and p-value < 0.001 for YG; in gamma band for 20%-40% of the GC: p-value < 0.001 both for EG and YG); in gamma band for 70%-90% of the GC: p-value = 0.046 for EG and p-value = 0.003 for YG). In all cases TW displayed higher coherence compared to SW. To summarize, each corticomuscular couple presents a significant difference between tasks in at least one time-frequency portion and in each group. In portions that present significant differences between tasks, the volume above threshold follows a particular trend both for YG and EG. In the EG, volume above threshold during DT is higher with respect to SW, as well as volume during TW is increased with respect to SW. Instead, the YG presents similar volume during SW and DT, while they show increased volume during TW compared to SW (fig. 3). Inter/intramuscular couples present in at least three time-frequency portions a significant task or interaction effect. In portions that present significant differences between tasks, the volume above threshold follows a particular trend similar for YG and EG: the volume is similar during DT and SW, while it is increased during TW compared to SW. In general, the increase of coherence from SW to TW is higher in YG compared to EG (fig. 3).

Concerning the stride duration analysis (table 3 and fig. 4) Friedman test detected a significant task effect for both YG (p-value < 0.001) and EG (p-value = 0.013). Post-hoc analysis revealed significant difference in median stride duration between SW and DT only for EG (p-value = 0.017). The median stride duration was higher in DT. Both YG (p-value < 0.001) and EG (p-value = 0.01) showed significantly different median stride duration during SW and TW. The median stride duration was higher in TW for both groups, even if the increase was more marked in YG. DT and TW showed significant difference in the median stride duration only for YG (p-value < 0.001). The median stride duration was higher in TW. Mann-Whitney U test detected a significant group effect only in TW (p-value = 0.001). The median stride duration was higher in YG compared to EG.

**Table 3.** Median (first quartile -third quartile) of stride duration for the two groups of subjects during the different walking conditions.

| Group | Stride duration: median (first quartile - third quartile) |  |  |
| --- | --- | --- | --- |
|  | SW | DT | TW |
| EG | 1.52 (1.41 - 1.72) | 1.75 (1.51 - 1.89) | 1.78 (1.68 - 1.86) |
| YG | 1.48 (1.37 - 1.64) | 1.65 (1.55 - 1.80) | 2.16 (1.94 - 2.32) |

**Figure 4.**
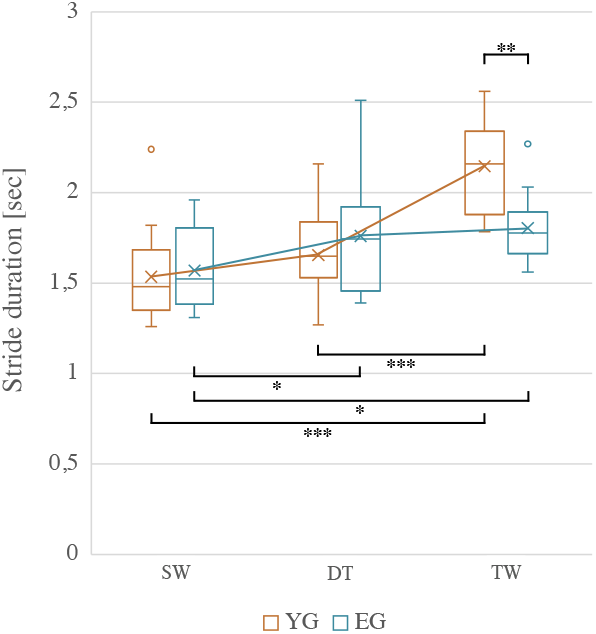
Stride duration for the two groups of subjects (EG = elderly group and YG = young group) during the different walking conditions (SW = spontaneous walking, DT = dual-task walking and TW = targeted walking).

## 4. Discussion

EEG and EMG are widespread and well-known quantitative techniques used for gathering biological signals at cortical and muscular level, respectively. Very recently, their combined use started to be considered as a potential breakthrough approach for neuromotor integrity/impairment analysis, evaluation, and assessment.

In this work, we exploited EEG and EMG to compute the corticomuscular, intermuscular, and intramuscular coherences of healthy young people and healthy elderly that walked in three conditions: i) spontaneous walking, dual-task walking, and iii) targeted walking. These walking conditions were chosen on the hypothesis that they progressively shift the subject’s cognitive functions (i.e., the CR) from the domain of cognitive activities (e.g., mathematical task), to the domain of motor activities (e.g., walking with precise targets to be followed). We focused on distal leg muscles (i.e., soleus and tibialis anterior of both legs), and the right and left hemisphere of the motor cortex (i.e., C3 and C4 EEG electrodes). Only single support phases (i.e., 20%-40% and 70%-90%) of the gait cycle were retained to avoid influences of possible motion artifacts on EEG signals during heel strikes, and beta and gamma frequency bands were analyzed separately.

For each couple of channels under investigation, we detected a significant difference between tasks at least in one phase of the gait cycle and frequency band, supporting the hypothesis that corticomuscular, intermuscular, and intramuscular coherences are able to reflect the modulation of corticomuscular communication between brain and muscles according to the cognitive or motor demand of the task. This is in line with previous results in literature finding coherence modulation with the task [2, 5, 7].

Comparing spontaneous and dual-task walking it can be observed that:

1. In young adults, corticomuscular coherence for all EEG-EMG couples considered did not significantly differ between spontaneous and dual-task walking, while it increased from spontaneous to dual-task walking in elderly;
2. Inter/intramuscular coherence for all EMG-EMG couples considered did not significantly differ between spontaneous and dual-task walking for both young and elderly participants;
3. Median stride duration increased for both young and elderly in dual-task compared to spontaneous walking.

The absence of significant differences in corticomuscular and inter/intramuscular coherence between spontaneous and dual-task walking in young participants confirmed our hypothesis that, in physiological conditions, gait is not a demanding task that requires conscious attention for young people. This suggests that they have an extended CR that can be used to perform more complex cognitive tasks during gait. This is in line with results by Filli et al. [2] that observed few changes in tibialis anterior and medial gastrocnemius activity between dual-task and regular walking in a healthy adult population (48.9±9.6 years old). Instead, the elderly showed an increase of corticomuscular coherence from spontaneous to dual-task walking, but the same increase was not observed in inter/intramuscular couples. This suggests that the elderly, when stressed at a cognitive level (e.g., when are asked to perform calculations while walking), make a greater cognitive effort that requires higher participation of the cortex (as explained by the corticomuscular coherence increase) compared to younger people. However, this increased level of concentration is not translated into an improvement of synchronous recruitment of distal leg muscles and gait performance. Indeed, the coherence between muscles did not increase. This can be hypothesized to be the effect of a reduced CR in elderly since they have been observed to require a greater cognitive effort compared to young people to perform the same dual-task. Due to limited CR, in elderly, the cognitive task (i.e., in our case calculations) probably engages subcortical regions that should normally be involved in gait control, as supported by the increment of corticomuscular coherence. On the contrary, younger people that have a greater CR, i.e., a greater neural capacity that results in more flexible and efficient cognitive and motor strategies, are able to compensate for the additional cognitive load imposed probably via the recruitment of additional brain regions or exploitation of other neural circuits not involved in gait control. Comparing spontaneous and targeted walking it can be observed that:

1. Corticomuscular coherence in targeted walking increased for all EEG-EMG couples considered compared to spontaneous walking both in young and elderly participants;
2. Inter/intramuscular coherence in targeted walking increased for all EMG-EMG couples compared to spontaneous walking both in young and elderly participants. The increase was more marked in young compared to the elderly group;
3. Median stride duration increased for both young and elderly in targeted compared to spontaneous walking, but the increase was more marked in young.

The increase of corticomuscular and inter/intramuscular coherence that was observed in targeted walking compared to spontaneous walking both in young and elderly adults is in line with previous results by Spedden et al. [5] that found greater corticomuscular and tibialis anterior intramuscular coherence during visually-guided walking compared to spontaneous walking. The increase of both corticomuscular and inter/intramuscular coherence between spontaneous and targeted walking in the young group suggests that young adults, when asked to perform a motor task that is guided by visual control, are able to transfer the increased cognitive effort (higher corticomuscular coherence) in a more fine tuning of the distal leg coordination (higher inter/intramuscular coherence) to achieve very precise foot placement. This supports the hypothesis that young adults have an extended CR that can be used to perform not only more complex cognitive functions but also more complex motor tasks. The minor increase in inter/intramuscular coherence in targeted walking that was observed in the elderly suggests that they have minor capability to integrate visuomotor components to coordinate distal leg muscles during gait when asked to perform precision motor tasks. These results are confirmed by the fact that the increase in stride duration from spontaneous to targeted walking was higher in the young group compared to the elderly group. Indeed, young adults reduce gait speed to exert more fine motor control and achieve better motor performance in term of precision of foot placement.

## 5. Conclusions and future developments

This study explored corticomuscular, intermuscular, and intramuscular coherences in healthy young people and healthy elderly during three different walking conditions that were chosen to place varying demands on the neuromuscular system. We observed higher corticomuscular and inter/intramuscular coherences during targeted walking compared to spontaneous walking in both groups, even if the increase was greater in young people. Considering dual-task walking compared to spontaneous walking, only corticomuscular coherence in the elderly increased. These results suggest age-related differences in cognitive reserve that reflect different abilities to perform complex cognitive or motor tasks during gait. We therefore demonstrated the feasibility and effectiveness of the proposed method to:

1. 1. investigate brain-to-muscle connectivity during different gait conditions, providing insights into the neural mechanisms underlying motor control during locomotion;
2. 2. investigate the functional changes that occur with aging in neuromuscular control of gait;
3. 3. quantify the cognitive reserve.

This methodology could be proposed as a novel clinically reliable tool to evaluate gait function and quantify the cognitive reserve. Future studies could extend the analysis to people with damages at the central nervous system affecting gait function, to explore the functional changes that occur with pathology in the neuromuscular system, and trace changes in corticospinal drive as induced by neurorehabilitation interventions or during disease progression. Moreover, the analysis could be extended also to proximal leg muscles and other motor and premotor brain source locations.

## Supporting information

Supplementary materials including 1 table and 3 figures

## 6. Acknowledgements

The authors thank the volunteers that participated in the study.

