## Supplementary materials including 1 table and 3 figures for "Brain-muscle connectivity during gait: corticomuscular coherence as quantification of the cognitive reserve"

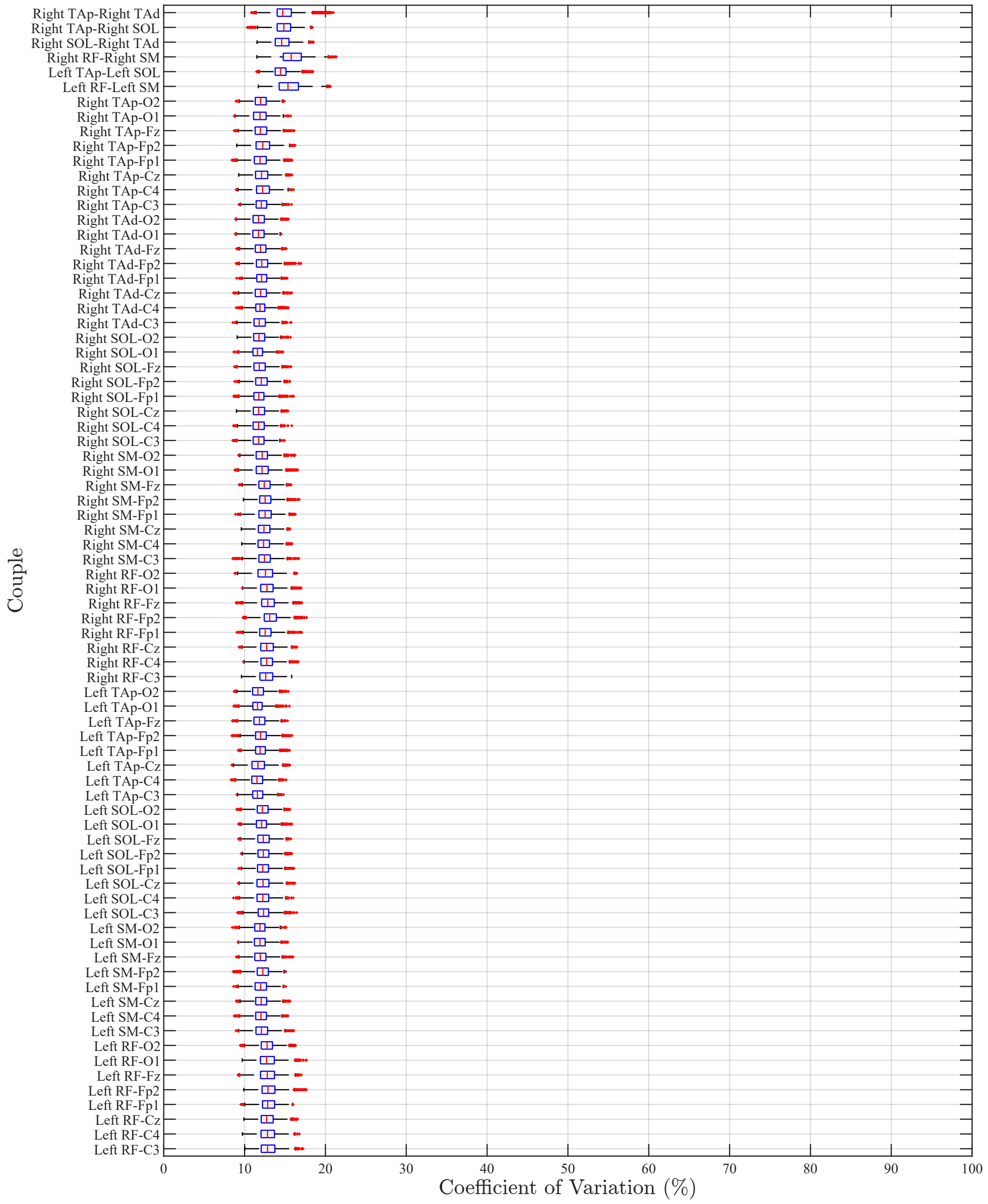


Figure 1: Coefficient of variations for each couple of channels in 15 - 60 Hz range. The first six are couples of two different muscles, the remaining are brain source location - muscle couples.

| Couple | Group | Task | Volumes above threshold: median (first quartile - third quartile) | | | |
|---|---|---|---|---|---|---|
| | | | 20% - 40% of GC<br>$\beta$ frequency band | 70% - 90% of GC<br>$\beta$ frequency band | 20% - 40% of GC<br>$\gamma$ frequency band | 70% - 90% of GC<br>$\gamma$ frequency band |
| Left SOL - C4 | EG | SW | 21.5 (15.9 - 28.1) | 24.6 (17.1 - 32) | 55.9 (43.7 - 74.8) | 66.2 (53.7 - 73.2) |
| | | DT | 31.8 (22.2 - 47.9) | 35 (30.8 - 56.5) | 52.2 (24.3 - 75.4) | 60.9 (43.5 - 73.7) |
| | | TW | 22 (13.2 - 35.2) | 51.2 (25.3 - 55.4) | 50.3 (41.9 - 58.3) | 70.8 (44.1 - 77.7) |
| | YG | SW | 15.6 (10.6 - 32.1) | 28.4 (13.1 - 41) | 36.4 (29.3 - 73.6) | 48.8 (32.9 - 81.4) |
| | | DT | 20.3 (9.3 - 33.3) | 20.4 (15.8 - 23.9) | 56.1 (31.7 - 70) | 40.9 (34.4 - 50.8) |
| | | TW | 23.6 (14.4 - 35) | 37.6 (34.4 - 51.3) | 47.3 (28.5 - 60.4) | 68.3 (54.7 - 87.6) |
| Left TAp - C4 | EG | SW | 29.2 (25.5 - 35.6) | 24.4 (20.2 - 28.4) | 74.4 (49.4 - 77.9) | 54 (43.3 - 61.9) |
| | | DT | 44.3 (41 - 52.5) | 37.7 (31.3 - 48.2) | 56.7 (47.1 - 64.8) | 64.4 (55.3 - 70.4) |
| | | TW | 51.4 (41.5 - 59.6) | 42.3 (32 - 60.9) | 68.5 (55.6 - 79) | 69.1 (50.4 - 78.6) |
| | YG | SW | 28 (18.7 - 57.8) | 11.5 (6.7 - 22.5) | 55 (38 - 71.3) | 34.7 (27.9 - 48.9) |
| | | DT | 34.3 (19.6 - 38.8) | 12.9 (3.6 - 21.9) | 59.4 (45.7 - 67.3) | 47.2 (37.4 - 60.3) |
| | | TW | 40.3 (37.4 - 56.6) | 23.2 (12.2 - 34.2) | 67.7 (51.1 - 82.3) | 55.3 (46.3 - 69.7) |
| Left SOL - Left TAp | EG | SW | 53.7 (45.2 - 59.4) | 37.1 (24.8 - 51.6) | 81.7 (67.1 - 97.8) | 77 (42 - 105.5) |
| | | DT | 55.7 (45.4 - 65.6) | 36.6 (24.9 - 56) | 101.3 (69.9 - 114.1) | 64.2 (38 - 72.7) |
| | | TW | 43.7 (27 - 56.5) | 69.8 (43.5 - 84.2) | 84.1 (62.6 - 98.7) | 62 (53.4 - 94.2) |
| | YG | SW | 59.7 (43 - 69.2) | 57.4 (28.6 - 84.2) | 91 (57.8 - 121.9) | 71.1 (48 - 112.9) |
| | | DT | 46.6 (41 - 61.2) | 60 (29 - 92.3) | 97 (66.7 - 124.6) | 87.8 (60.7 - 149.7) |
| | | TW | 69.8 (44.1 - 92.9) | 82.7 (75.4 - 88.8) | 126.8 (83.9 - 151.4) | 114 (92.5 - 125.7) |
| Right SOL - C3 | EG | SW | 32.1 (25.1 - 41.5) | 18.2 (8.4 - 27.2) | 49.9 (38.6 - 70.1) | 35.5 (31.8 - 44.8) |
| | | DT | 32.7 (20.1 - 53) | 18.7 (14.3 - 28.1) | 58.1 (44.5 - 79) | 50.4 (43.1 - 74.6) |
| | | TW | 39.6 (35.4 - 51.7) | 18.1 (13.3 - 26.5) | 66.5 (57.4 - 75.6) | 59.3 (49.8 - 66.1) |
| | YG | SW | 22.4 (10.2 - 49.5) | 20.7 (9.1 - 26.7) | 42.8 (35.1 - 72.4) | 35.4 (27.1 - 46.1) |
| | | DT | 33.1 (17 - 49.6) | 14 (11 - 31.6) | 54.2 (40.7 - 73.3) | 42.6 (30.8 - 82.1) |
| | | TW | 57.8 (49.1 - 59.7) | 25.1 (14 - 42.9) | 72.9 (66.6 - 80.5) | 53.5 (39.1 - 70) |
| Right TAp - C3 | EG | SW | 17.2 (10.2 - 26.5) | 32.2 (26.4 - 33.9) | 53.9 (39.8 - 62.6) | 50.6 (26.9 - 65.6) |
| | | DT | 36.4 (29.7 - 47.6) | 38.3 (30.7 - 50.2) | 65.8 (43.6 - 68) | 61.6 (36.6 - 75.8) |
| | | TW | 35.2 (14.7 - 46.8) | 33.1 (27.2 - 46.3) | 66.5 (43.7 - 78) | 66.6 (61.3 - 85.3) |
| | YG | SW | 10.3 (3.1 - 24.8) | 24.8 (15 - 42.5) | 50.2 (19 - 64.7) | 51.4 (24.5 - 64) |
| | | DT | 9.4 (7 - 24.9) | 32.2 (27 - 36.4) | 59.9 (39.4 - 70.1) | 62.3 (43 - 76.2) |
| | | TW | 33 (14.2 - 43.7) | 33 (18.4 - 48.2) | 80.2 (60.5 - 87) | 64.5 (35 - 87.2) |
| Right SOL - Right TAp | EG | SW | 31.4 (22.6 - 42.9) | 36.7 (28.7 - 60) | 59 (41.8 - 81.2) | 73.8 (42.2 - 104.1) |
| | | DT | 37.8 (17 - 54.1) | 58.1 (41.2 - 66.2) | 75.8 (59.3 - 84.5) | 74.8 (52.6 - 92.4) |
| | | TW | 66.3 (41.4 - 80.3) | 53.5 (50.5 - 56.7) | 92.6 (79.8 - 112.4) | 107.7 (79.2 - 114.9) |
| | YG | SW | 56.1 (33.6 - 76.1) | 59.5 (48.1 - 79.8) | 101.9 (80.9 - 129) | 105.3 (81.4 - 134) |
| | | DT | 51.2 (36.9 - 79.4) | 65.2 (45.5 - 107) | 99 (80.4 - 130.6) | 108.8 (86.5 - 166.3) |
| | | TW | 82.2 (66.7 - 98.8) | 103.4 (72.1 - 112.9) | 106 (94.1 - 131.1) | 147.9 (115 - 157.3) |
| Right TAp - Right TAd | EG | SW | 120.5 (112.5 - 132.2) | 105.7 (95.3 - 114.8) | 159.7 (157.9 - 170.1) | 141.1 (129.9 - 159.4) |
| | | DT | 131.5 (126.3 - 135.3) | 104.6 (100.6 - 117) | 171.6 (160.1 - 180.4) | 144.8 (135.8 - 149.6) |
| | | TW | 145.6 (139.3 - 161.3) | 116.5 (108.9 - 123.9) | 196.1 (187 - 215.6) | 158 (147.7 - 168.5) |
| | YG | SW | 125.3 (108.7 - 138.8) | 121.7 (109.2 - 126.9) | 160.1 (140.1 - 190.3) | 166.9 (139.6 - 170.4) |
| | | DT | 135.3 (92.7 - 156) | 123.6 (111.8 - 132.1) | 184.3 (136 - 199.1) | 165.9 (147.9 - 177.3) |
| | | TW | 175 (158.2 - 185.9) | 128.6 (123.8 - 138.2) | 231.7 (204.3 - 251.1) | 169 (157.5 - 182.9) |

Table 1: Median (first quartile - third quartile) of the volumes above threshold for corticomuscolar coherences involving SOL and TAp muscles of both legs and the correspondent controlateral EEG locations on the central region (C3, located on the left, and C4, located on the right), for the intermuscular coherences between SOL and TAp of legs and for the intramuscular coherence between TAp and TAd of the right leg, for the two groups of subjects (EG = elderly group and YG = young group) during the different walking conditions (SW = spontaneous walking, DT = dual-task walking and TW = targeted walking). Volume above threshold is measured in $Hz \cdot \%GC$. GC = gait cycle.

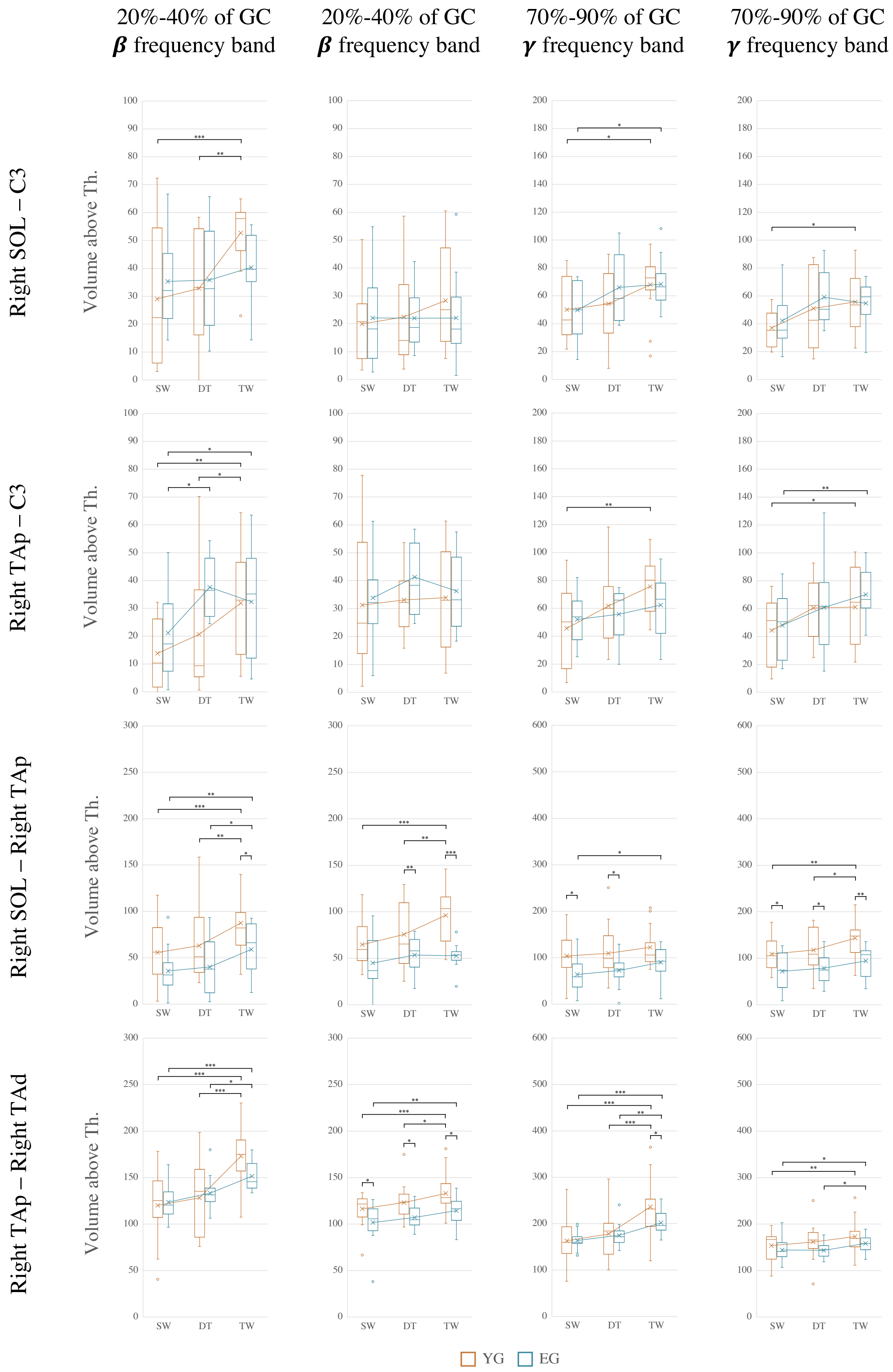


Figure 2: Distribution of the volume above threshold of corticomuscular and inter/intramuscular couples involving right muscles. Each row represents a couple of channels and each column a time-frequency portion of coherence. Each figure presents the distribution of the volume above threshold for the two groups (orange for the elderly group - EG - and light blue for the young group - YG), and the three tasks (SW = spontaneous walking, DT = dual-task walking and TW = targeted walking). The cross of each boxplot represents the mean of the distribution. Statistical significance of the comparisons between tasks and groups is displayed by asterisks. Volume above threshold is measured in $Hz \cdot \%GC$. GC = gait cycle.

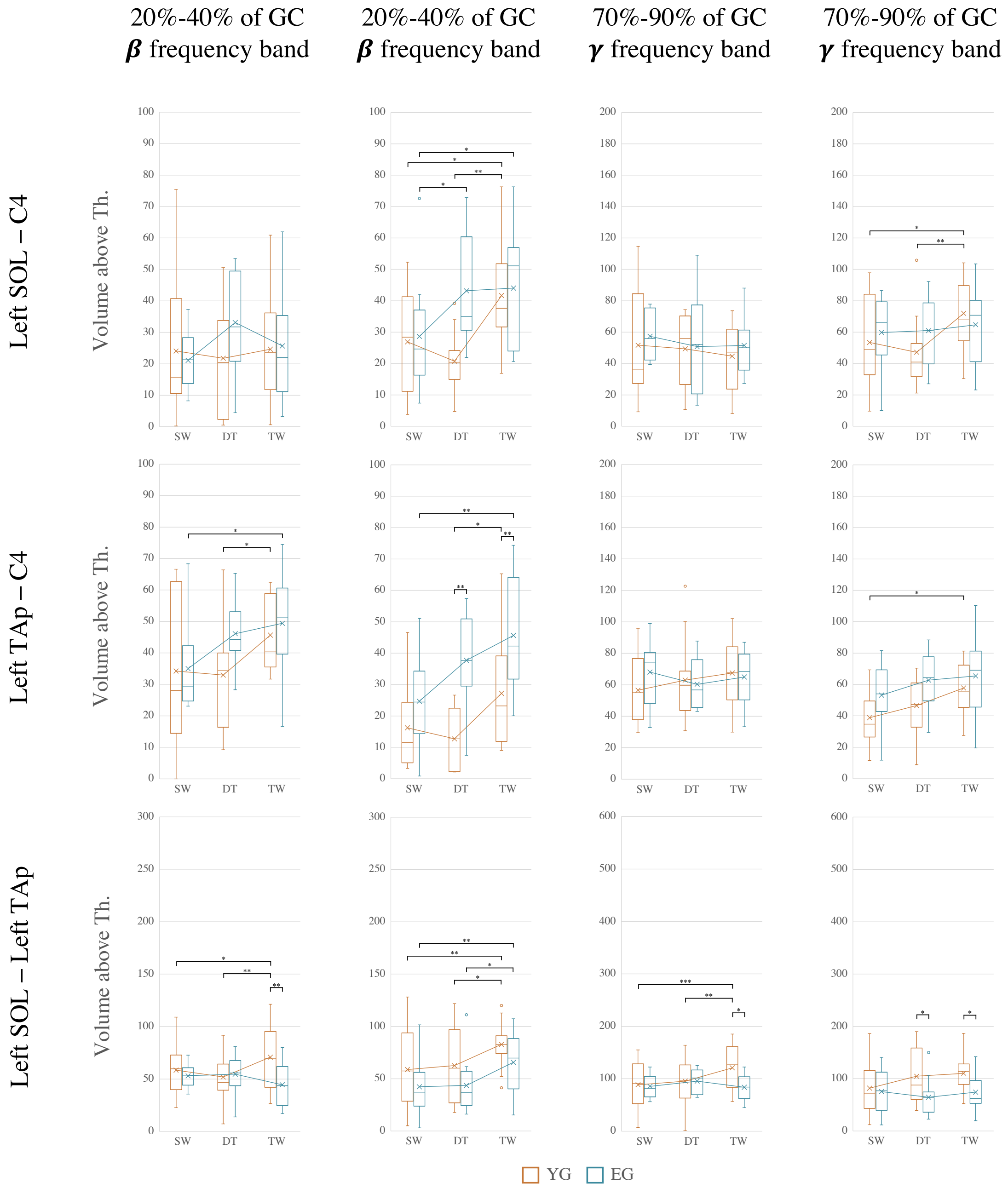


Figure 3: Distribution of the volume above threshold of corticomuscular and intermuscular couples involving left muscles. Each row represents a couple of channels and each column a time-frequency portion of coherence. Each figure presents the distribution of the volume above threshold for the two groups (orange for the elderly group - EG - and light blue for the young group - YG), and the three tasks (SW = spontaneous walking, DT = dual-task walking and TW = targeted walking). The cross of each boxplot represents the mean of the distribution. Statistical significance of the comparisons between tasks and groups is displayed by asterisks. Volume above threshold is measured in $Hz \cdot \%GC$. GC = gait cycle.
